# Differential impact of endogenous and exogenous attention on activity in human visual cortex

**DOI:** 10.1101/414508

**Authors:** Laura Dugué, Elisha P. Merriam, David J. Heeger, Marisa Carrasco

**Author notes:** **Corresponding author:** Laura Dugué, Current address: 45 rue des Saints-Pères 75006 Paris, FRANCE.

## Abstract

How do endogenous (voluntary) and exogenous (involuntary) attention modulate activity in visual cortex? Using ROI-based fMRI analysis, we measured fMRI activity for valid and invalid trials (target at cued/un-cued location, respectively), pre- or post-cueing endogenous or exogenous attention, while participants performed the same orientation discrimination task. We found stronger modulation in contralateral than ipsilateral visual regions, and higher activity in valid-than invalid-trials. For endogenous attention, modulation of stimulus-evoked activity due to a pre-cue increased along the visual hierarchy, but was constant due to a post-cue. For exogenous attention, modulation of stimulus-evoked activity due to a pre-cue was constant along the visual hierarchy, but was not modulated due to a post-cue. These findings reveal that endogenous and exogenous attention distinctly modulate activity in visuo-occipital areas during orienting and reorienting; endogenous attention facilitates both the encoding and the readout of visual information whereas exogenous attention only facilitates the encoding of information.

## INTRODUCTION

Spatial, covert visual attention is the selective processing of visual information in space, without change in gaze. Attention can be allocated voluntarily –endogenous attention– or involuntarily –exogenous attention. Endogenous and exogenous attention have different temporal dynamics; endogenous takes about 300 ms to be deployed and can be sustained at will whereas exogenous attention takes only about 100 ms to be deployed and it is transient (for reviews see ^1,2^). These two types of attention often have similar perceptual consequences (reviews by ^1,2^), but notable exceptions indicate that whereas endogenous attention acts in a flexible way, exogenous attention acts in an inflexible or automatic way. For instance: (a) The benefits and costs in perception (discriminability and speed of information accrual) scale with cue validity for endogenous but not for exogenous attention (e.g. ^3^); (b) The effects of covert attention on contrast sensitivity often differ for endogenous and exogenous attention (e.g. ^4–9^); and (c) For a texture segmentation task in which heightened spatial resolution improves or impairs performance as a function of target eccentricity, endogenous attention improves performance across eccentricity, whereas exogenous attention improves performance at peripheral locations where resolution is poor, it hampers performance where resolution is already high for the task at hand (e.g. ^10–15^).

Basic visual processes, such as contrast sensitivity and spatial resolution, are mediated by activity in early visual cortex, and are altered by covert attention (reviews by ^1,2,16,17^). Specifically, single-unit studies in monkeys have demonstrated effects of endogenous attention in occipital areas (e.g. ^18–25^). Additionally, fMRI studies have shown that endogenous attention causes a baseline shift in early visual areas (e.g. ^26–32^; review by ^33^) and increases the dynamic range of fMRI responses ^34,35^. Comparatively, little is known about the effect of exogenous attention on visual areas both from single-unit studies ^36,37^ and fMRI studies (for reviews see ^1,16^).

Since Corbetta and Schulman’s seminal review (2002) ^38^ on the neural bases of endogenous and exogenous attention, there has been emphasis on characterizing networks of brain regions within the frontal and parietal lobes (for reviews see ^33,39,40^). Yet, there remains considerable debate regarding the dissociation between dorsal regions, for endogenous attention, and ventral regions, for exogenous attention (e.g. ^40,41^), including the role of the temporo-parietal junction (TPJ; ^42–45^). Classically, researchers have described endogenous attention as a top-down process, and exogenous attention as a bottom-up process ^1,38–40,46–48^. This characterization originated in psychophysics experiments, and was then studied using fMRI, in which the two types of attention have been often investigated separately (for reviews ^1,33,39,46^).

Studies of endogenous and exogenous attention focusing on parietal and frontal areas have shown that the two types of attention differentially modulate fronto-parietal connectivity ^40,49^. For example, there are critical differences in the temporal order of neural responses in frontal and parietal cortex in monkeys between these attention conditions, i.e. frontal activity precedes parietal activity during endogenous attentional orienting, whereas parietal activity precedes frontal activity during exogenous orienting. Critically, it is often assumed that the effects of endogenous and exogenous attention are the same in striate and extra-striate areas^33,38,39,50,51^.

A number of important considerations limit the conclusions that may be drawn from the few studies that have directly compared independent effects of visual exogenous and endogenous spatial attention ^40,51–53^:

1. The effect of attention on behavioral performance was assessed in some studies with a detection task using RT as their only dependent variable ^51,53^, in which performance may differ due to speed of processing, discriminability or criterion ^54–56^ and/or motor behavior^57,58^.
2. In the studies in which performance was assessed in a discrimination task using RT ^52^, small RT differences were reported, which could have resulted from speed of processing, discriminability, or criterion factors ^54–56^.
3. In the studies in which accuracy was not assessed ^40,51,53^, it is not possible to know whether task difficulty was the same for both types of attention, and task difficulty can interact with the strength of fMRI activity ^59,60^.
4. For the studies in which eye position was not monitored while participants performed the task in the scanner ^40,51–53^, the results could be due to covert attention, overt attention or both ^1,61,62^.
5. Given that exogenous attention is a fast, transient process (for reviews see ^1,2,16,17^), it was inappropriately manipulated in the studies in which long stimulus onset asynchronies (SOA) were used ^51,52,63–65^ making the comparison between the two attention conditions problematic.
6. Except for two studies, one interested in attentional modulation in the Fusiform Face Area (FFA) ^65^ and the other in the TPJ ^42^, statistical parametric mapping was applied to group averaged data to identify regions of the brain that were active during task performance. Some found significant activity in the occipital pole ^51–53^, but attentional modulation of fMRI activity was not systematically assessed across different visual areas.

Given all these methodological limitations, it is unknown how these two types of attention affect neural activity in each individual visuo-occipital area, and how modulation of activity in visual cortex is linked to changes in perceptual performance. (See Dugué and colleagues ^42^, who published a table summarizing these and other methodological problems for studies regarding covert attention and TPJ activation).

Typically, covert attention is manipulated by presenting a pre-cue, prior to the target – and the aforementioned studies also did so (see **Table 1**). However, endogenous post-cues, presented after target offset, can also improve performance by affecting the information readout ^66–69^ and modulate fMRI activity in early visual areas ^26,68–70^. Exogenous post-cues also affect performance in some tasks ^71,72^, but not in others ^73–76^, and the only fMRI study evaluating post-cues in exogenous attention in occipital cortex showed no such modulation ^77^. Critically, no single study has compared visual cortex activity with post-cues in endogenous and exogenous attention.

**Table 1.**
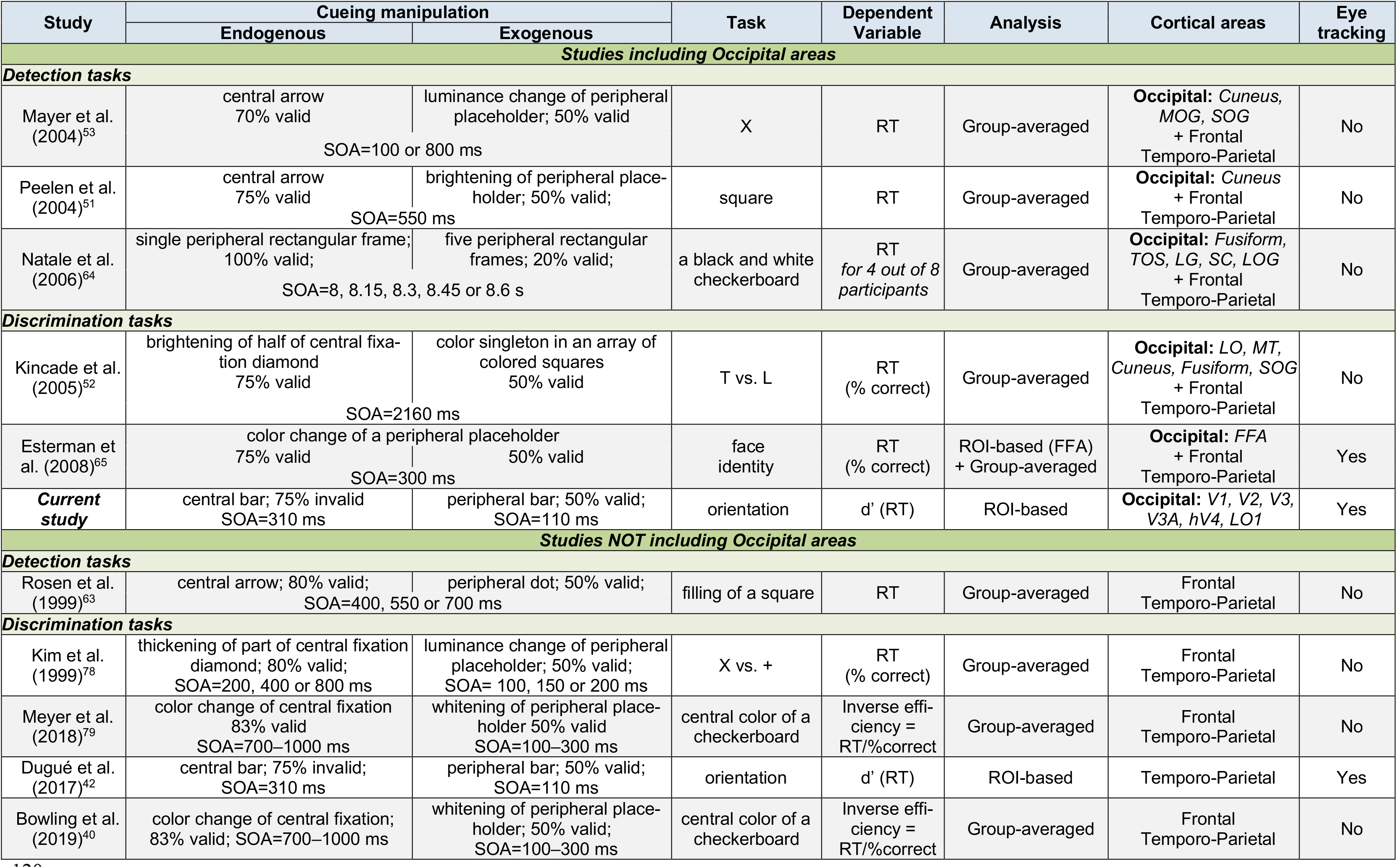
fMRI studies comparing endogenous and exogenous attention independently in human participants. For each study, we report the cueing manipulation for each attention condition (SOA: Stimulus Onset Asynchrony is the duration of the cue + interval before onset of the target), the task performed by the participants, the dependent variable reported in the publication, whether the analysis was based on group-averaging statistics or ROI-based on single-participants, the cortical areas reported in the publications and showing significant activation due to cue and/or target, and whether or not eye data were monitored in the scanner or used for the subsequent fMRI analysis. IOG: Inferior Occipital Gyrus; LG: Lingual Gyrus; LO: Lateral Occipital area; LOG: Lateral Occipital Gyrus; MOG: Middle Occipital Gyrus; MT: Middle Temporal area; SC: Striate Cortex; SOG: Superior Occipital Gyrus; TOS: Temporal-Occipital Sulcus.

Here, with the same participants, task, stimuli and task difficulty for all attention manipulations, we systematically tested the following four predictions: (1) Pre-cueing should induce an attentional modulation of fMRI activity (for reviews see ^1,16,33,39,40^), higher in the valid than the invalid condition in which attention needs to be reoriented to the opposite location (e.g. ^42,50,80,81^) to perform the task. (2) Both endogenous and exogenous pre- and post-cueing effects should be stronger in visual regions contralateral to the attended hemifield (e.g. ^26,77^). (3) Pre-cueing endogenous attention, but not exogenous attention, should increase activity modulations along the visual hierarchy (e.g. higher in V4 than in V1; ^26,31^; for reviews ^33,39^). For endogenous attention, a top-down process, modulations from higher-order, fronto-parietal attentional regions would send feedback information to visual cortex with diminishing effects in earlier visual areas ^31,32,39,82–85^. (4) Post-cueing endogenous ^26,68–70^, but not exogenous ^77^, attention should induce attentional modulation of fMRI activity in early visual areas. Voluntary, endogenous attention would facilitate reading out perceptual information ^42^, and modulate its processing ^26,68–70^. The only fMRI study assessing the effects of post-cueing exogenous attention in occipital cortex found no attentional modulation of fMRI activity ^77^; some behavioral studies report post-cueing effects ^71,72^ but others found no such effects ^73–76^.

To test these four predictions, we measured fMRI activity and compared the effects of endogenous and exogenous attention in early (V1, V2, V3) and intermediate (V3A, hV4, LO1) visual areas while the same participants performed the same task –a two-Alternative Forced Choice (2-AFC) orientation discrimination task, contingent upon contrast sensitivity ^4,5,8,80,86–89^. We used a fully-crossed design: two attention conditions –endogenous or exogenous attentional orienting– and two types of cueing –pre- or post-cue. We evaluated fMRI activity at both the attended and the un-attended locations, given the ubiquitous performance tradeoffs at attended (benefits) and unattended (costs) locations compared to a neutral condition (e.g. ^3,28,88–93^), and the importance of evaluating both the orienting and reorienting of attention, critical in an ever-changing environment ^42,80,81,94^.

This is the first study to systematically evaluate and directly compare how pre and post- orienting, and reorienting, of endogenous and exogenous attention modulate neural activity in visual cortex to affect behavior. The results indicate that these two types of spatial covert attention distinctly modulate activity in individual retinotopic visual cortical areas. These differences in activity are consonant with differential engagement of top-down and bottom-up processes and their respective temporal dynamics. These results suggest that endogenous attention facilitates both the encoding and the readout of visual information whereas exogenous attention only facilitates the encoding of information.

## RESULTS

### Endogenous and exogenous attention improve performance

Participants performed a 2-AFC orientation-discrimination task under two attentional conditions (exogenous or endogenous attention), when the cue was presented either before (precue) or after (post-cue) the grating stimuli (see Methods), while their brain activity was measured with fMRI (**Figure 1**).

**Figure 1.**
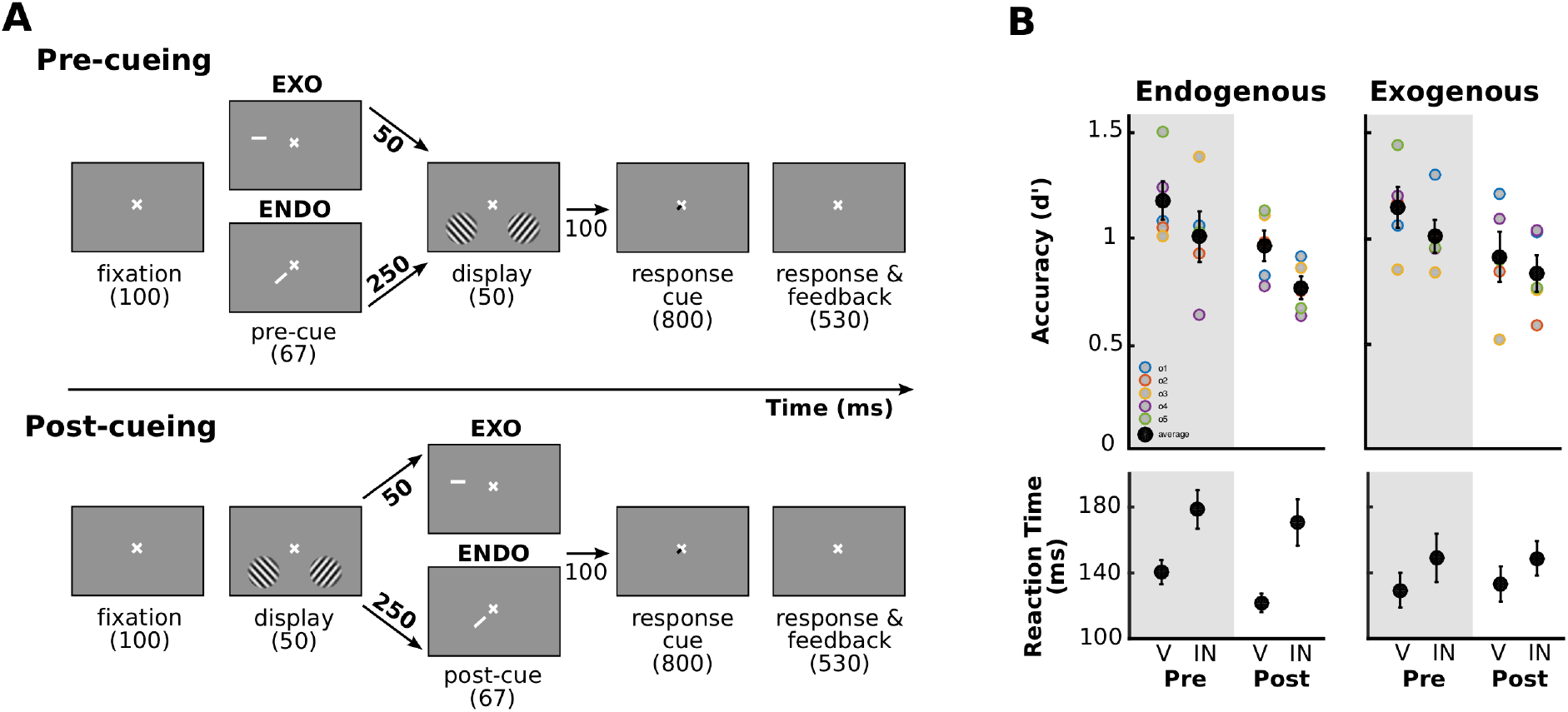
Experimental protocol. **A.** Participants performed a 2-AFC orientation-discrimination task. Pre-cues and post-cues were presented before and after the stimuli, respectively. Exogenous cues appeared in the periphery, above one of the two stimulus locations. Endogenous cues appeared at the center of the screen (the display is not at scale for visibility purposes) and indicated one of the two stimulus locations The ISI between the cue and the gratings was shorter for the exogenous (50 ms) than the endogenous (250 ms) conditions. A response cue indicated the target location and instructed participants to indicate whether the target grating was tilted clockwise or counterclockwise of vertical by pressing one of two keys. To provide feedback, the fixation-cross turned green or red for a correct or an incorrect answer, respectively. **B.** Behavioral performance averaged across participants (black dots) and for each of them (colored dots; n=5) for endogenous (left) and exogenous (right) attention. (Top) Performance accuracy (d’; top) and median reaction time (bottom) as a function of cueing condition. V, valid cue (same location of pre-cue/post-cue as response cue). IN, invalid cue (different location of pre-cue/post-cue than response cue). Pre, pre-cue presented before the stimuli. Post, post-cue presented after the stimuli. Valid cues induced more accurate and faster responses (there was no speed-accuracy trade-off). Error bars, ± 1 SEM across participants.

In each condition, we calculated performance accuracy (d’), as the main dependent variable, for each participant separately (**Figure 1B**, top row). A three-way repeated measures 2×2×2 ANOVA (exogenous/endogenous × valid/invalid × pre/post-cue) revealed higher performance for valid than invalid cues (F(1,4)=23.6, p=0.008), that the effect of exogenous and endogenous cues were statistically indistinguishable (F(1,4)<1), and that there was no significant difference between pre- and post-cues (F(1,4)<1). None of the two and three-way interactions were significant (F<1).

In each condition, we also calculated reaction time (RT), as a secondary dependent variable for each participant separately (**Figure 1B**, bottom row). Although the task is not optimal to asse RT effects due to the long response cue window during which participants are not allowed to provide their response, we assessed RT to rule out speed-accuracy tradeoffs. A three-way repeated measures ANOVA revealed faster reaction times for valid than invalid cues (F(1,4)=62.3, p=0.001). There was no significant difference between exogenous and endogenous cues (F(1,4)=2.7, p=0.17), or between pre- and post-cues (F(1,4)=1.7, p=0.27). Two significant interactions indicated that the differences between valid and invalid cues (F(1,4)=16.2, p=0.02) and between pre- and post-cues (F(1,4)=8.1, p=0.047) were more pronounced for endogenous attention than for exogenous attention.

These behavioral results, which are consistent with previous findings ^3–6,8,28,80,87–89,91^, show that attention improved orientation discrimination (d’ and reaction time), with no evidence of a speed-accuracy trade-off, and similarly for both types of attention and for both pre- and post-cues. Thus, the behavioral effects confirm the successful manipulation of endogenous and exogenous attention, for both pre- and post-cues, consistently across individuals.

### Attentional modulation of perceptual and post-perceptual information processing in visual cortex

Visual areas were mapped in each participant following retinotopic mapping procedures (**Figure 2**, left panel) and a targeted stimulus localizer (**Figure 2**, right panel), and regions of interest (ROIs) were selected based on previous literature: V1, V2, V3 (for V2 and V3, ventral and dorsal ROIs were averaged), V3A, hV4 and LO1 (e.g. ^95–98^).

**Figure 2.**
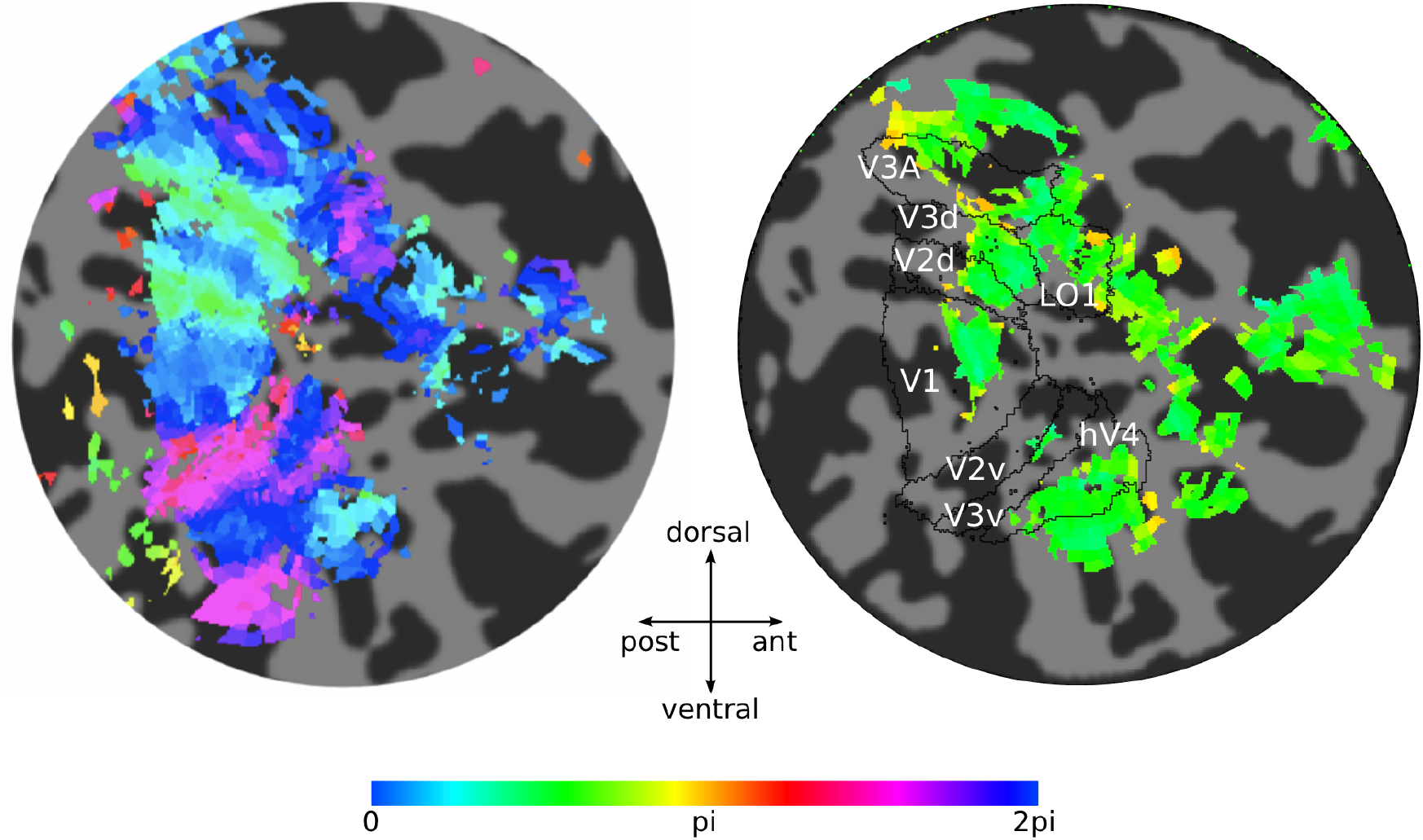
Retinotopic mapping and stimulus localizer of a representative participant. A flattened representation of the right hemisphere occipital pole. **Left**, map of polar angle. The color of each voxel in the map represents the response phase of the time series elicited by a rotating wedge stimulus. **Right**, stimulus localizer. The black outlines correspond to the retinotopic areas of interest (defined for each participant based on their polar angle maps). The color of each voxel indicates the phase of the response to the grating stimuli presented in the lower left visual field.

Activity was higher for contralateral than ipsilateral brain areas. For each ROI, we measured the fMRI response amplitudes for each type of attention – exogenous and endogenous – and each cueing condition – pre- and post-cueing, for the contralateral and ipsilateral side to the cued (attended) location. We analyzed the fMRI responses evoked by each type of attention in the contralateral and ipsilateral brain regions relative to the cue location (**Supplemental Figure 1**). ANOVAs indicated that there was higher contralateral than ipsilateral activity across brain areas (endogenous: F(1,4)=59.9, p=0.0015; exogenous: F(1,4)=218.8, p=0.0001). For both types of attention, this difference was more pronounced for valid than invalid cues (endogenous: F(1,4)=8.6, p=0.04; exogenous: F(1,4)=21.1, p=0.01). In the following analyses, we then concentrate on fMRI activity in the contralateral ROIs to the attended location. **Supplemental Figure 2** shows mean fMRI responses for a representative participant for each attention condition for two contralateral ROIs (V2 and LO1) relative to the cue. Response amplitude was obtained from the target areas indicated in **Figure 2** **(right panel)**.

We calculated an attentional modulation index: AMI = Vc / average (Vc + Ic + Vi + Ii), which takes into account the overall response magnitude of each ROI. The AMI corresponds to the ratio of the activity in contralateral ROIs for valid trials (Vc) over the average activity of contralateral valid (Vc) and invalid trials (Ic), and distractor stimuli for pre and post-cue trials for ipsilateral valid (Vi) and invalid trials (Ii). This AMI takes advantage of our fully factorial design. We are isolating the activity brought about by the relevant attentional condition (valid trials in endogenous or exogenous condition) and comparing it with the overall response magnitude elicited by the presence of the stimuli regardless of attentional states. (Because some condition values were negative, we added a fixed constant (0.2) to all condition values of each participant before computing the AMI so that all resulting values were positive; see original values in **Figure 5.** Below we show that the pattern of results is the same for different constant values, indicating that our results do not depend on the specific constant value we selected). A ratio of 1 would indicate no attentional modulation. The larger the AMI ratio, the higher the modulation in the valid trials. We note that the AMI that we report cannot be directly compared with the AMI or the values used by others, neither can the magnitude of the attentional modulation of previous studies can be directly compared. From the list of the studies we include in Table 1, most use group-average analyses, and statistics on contrasts between different conditions ^51–53,64,78,79^, and two used ROI-based analyses but did not use AMI ^42,65^. Other studies using ROI-based analyses plot the different conditions individually, or differences between conditions (e.g. ^32,70^), and yet others use AMI indices but they differ among studies (e.g. ^26,31,77,99^).

First, we showed that the AMI was significantly higher than 1 when combined across all conditions and ROIs (t(4) = 6.7, p = 0.0026, Cohen’s d = 2.99), and this was the case for each participant. These results are consistent with previous findings showing that attentional orienting increases fMRI signal in early visual cortex (for review ^33^).

We then tested our two novel predictions: (1) attentional modulation of fMRI stimulus-evoked activity increases along the visual hierarchy for endogenous pre-cueing, but is constant for exogenous pre-cueing; and (2) attentional modulation of fMRI stimulus-evoked activity for endogenous post-cueing is constant along the visual hierarchy, whereas there is no attentional modulation for exogenous post-cueing. We first calculated the differences in activity between valid and invalid trials for each type of attention and for pre- and post-cues, across the hierarchy of visual cortical areas. fMRI (AMI) responses were significantly larger for valid than invalid cues, for both endogenous and exogenous cues, and for both pre- and post-cues. A three-way repeated measures ANOVA (2 exogenous/endogenous × 2 pre/post-cue × 6 ROIs) of the difference between valid and invalid conditions revealed significant main effects of exogenous/endogenous condition (F(1,4) = 33.8, p = 0.0043) and of ROI (F(5,4) = 3.3, p = 0.0258), as well as an interaction between pre/post-cue and ROI (F(5,4) = 3.5, p = 0.0192). Thus, we investigated modulations of fMRI activity separately for pre and post-cueing attention conditions.

Linear mixed models ^100^ were then used to test: (1) an increase of fMRI activity along the visual hierarchy for pre-cueing endogenous attention but not for exogenous attention, and (2) post-cue attentional modulation of fMRI activity for post-cueing endogenous attention but not for exogenous attention. We implemented two models, one for the pre-cueing and the other for the post-cueing condition, to test the link between fMRI activity and the interaction between ROI and endogenous/exogenous attention conditions. In each model, we entered as fixed effect the ROI and the endogenous/exogenous conditions into the model, as well as their interaction. As random effects, we had participant’s intercepts and slopes for the effect of ROI and endogenous/exogenous conditions. Visual inspection of residual plots did not reveal any obvious deviations from homoscedasticity or normality. P-values were obtained by likelihood ratio tests. For the pre-cueing condition, we observed a significant effect of ROI (t(56)=3.5, p=0.0009, estimate=0.05±0.01, standard error), as well as a significant interaction between ROI and endogenous/exogenous conditions (t(56)=-2.3, p=0.025, estimate=-0.018±0.008, standard error). For the post-cueing condition, we observed a significant effect of the fixed effect intercept (t(56)=23.3, p<0.0001, estimate=11±0.05, standard error). Finally, we performed independent regression analyses on each pre-/post-cueing and endogenous/exogenous attention conditions (**Figure 3**). We observed a significant increase along the hierarchy of the visual areas of the activity difference evoked by valid and invalid trials for the endogenous pre-cueing condition (F(4)=16.5, p=0.015, R^2^=0.81), but not for the exogenous pre-cueing condition (F(4)=1.8, p=0.249, R^2^=0.31), endogenous post-cueing (F(4)=2.2, p=0.216, R^2^=0.35) or exogenous post-cueing conditions (F(4)=0.3, p=0.61, R^2^=0.07). See **Supplemental Figure 3** and **Supplemental Table 1 and 2**, which show the same pattern of results for different constant values, indicating that our results do not depend on the specific value of the constant we selected.

**Figure 3.**
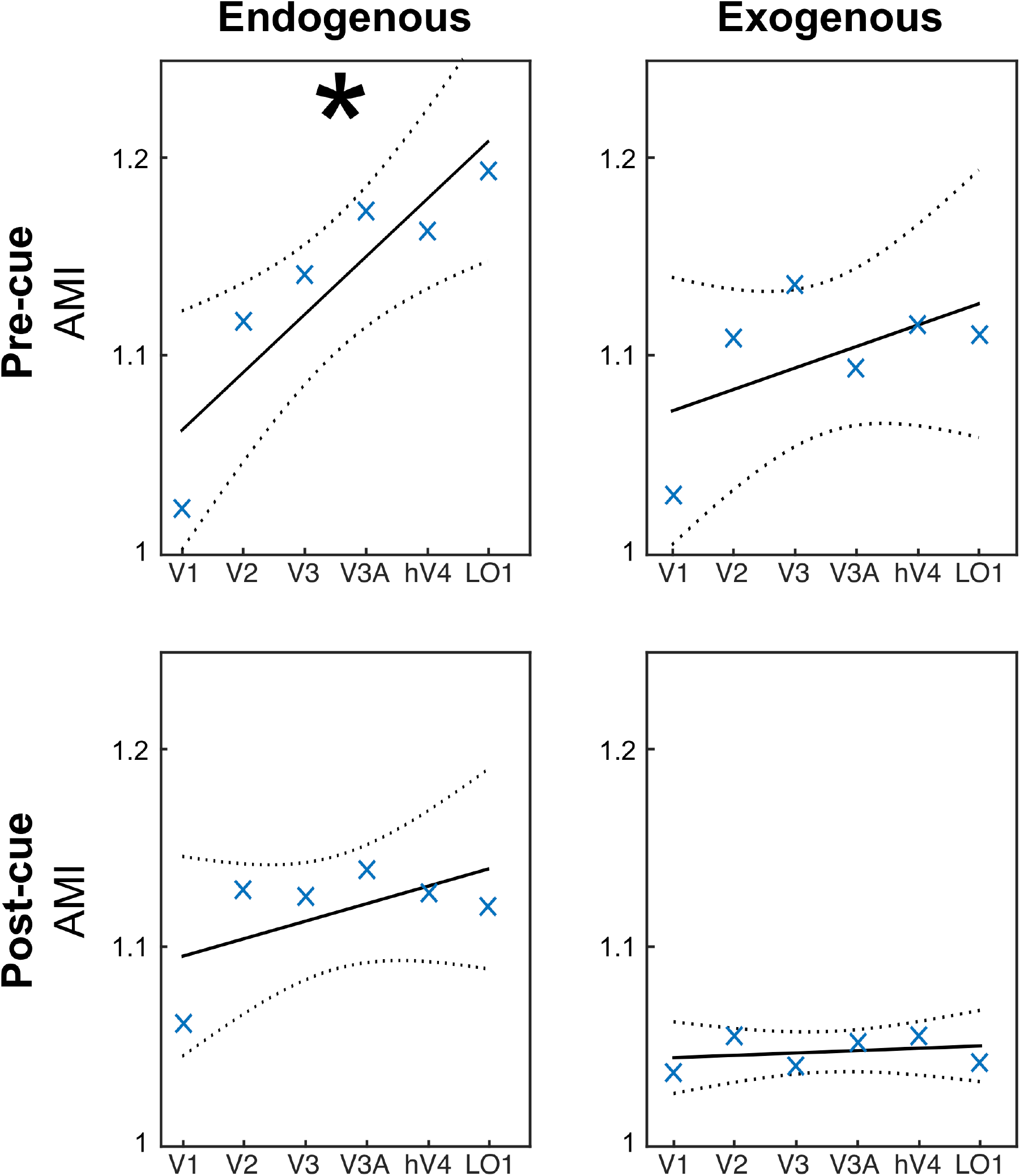
Specificity of pre and post-cueing for endogenous and exogenous attention. AMI, Attentional Modulation Index separately for pre- and post-cue conditions for each ROI: AMI = Vc / average (Vc + Ic + Vi + Ii). Vc, valid trials in contralateral ROI. Ic, invalid trials in contralateral ROI. Vi, valid pre- and post-cue trials in ipsilateral ROI. Ii, invalid pre- and post-cue trials in ipsilateral ROI. The larger this ratio, the higher the modulation in the valid trials. *, Statistically significant regression analysis (p < 0.05).

Together, the results of the linear mixed models and the regression analyses (**Figure 3**) showed differential effects between pre-/post-cueing and endogenous/exogenous attention conditions. The AMI increases along the hierarchy of the visual areas in pre-cueing of endogenous attention but not for pre-cueing of exogenous attention. Moreover, the AMI with post-cueing of endogenous attention is constant along the hierarchy (significant intercept), but absent for post-cueing of exogenous attention. These analyses are consistent with post-hoc t-tests performed for each condition and ROI in the AMI modulation (**Figure 4**). See **Supplemental Figure 4**, which shows the same pattern of results for the difference in fMRI activity (in percent signal change) between valid and invalid.

**Figure 4.**
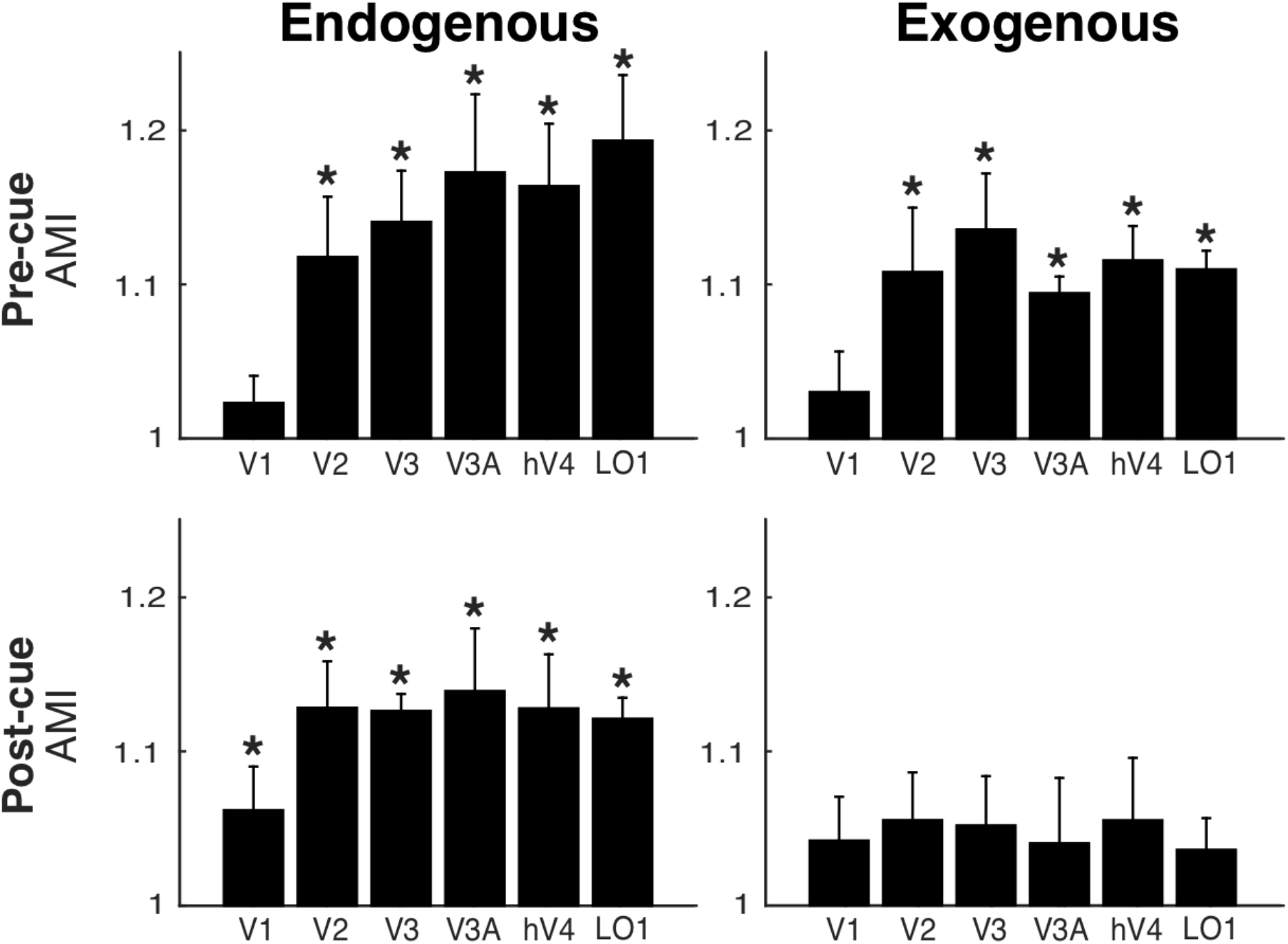
Single ROI responses for pre and post-cueing and for both endogenous and exogenous attention. The AMI is plotted separately for pre and post-cue conditions for each ROI (same data as in Figure 3). *, Statistically significant difference between valid and invalid, separately for pre and post-cueing (p < 0.05). Error bars on plots are ± 1 SEM.

For both endogenous attention (p < 0.05 for V2, V3, V3a, V4 and LO1; one-tailed t-tests) and exogenous attention (p < 0.05 for V2, V3, V3A, hV4 and LO1; one-tailed t-tests) pre-cues elicited greater fMRI activity for valid than invalid cues (**Figure 4**, top-left and top-right panels). Furthermore, for endogenous attention (p < 0.05 for V1, V2, V3, V3A, hV4 and LO1; one-tailed t-tests; **Figure 4**, bottom-left panel), but not for exogenous attention (all p > 0.05; one-tailed t-tests; **Figure 4**, bottom-right panel), post-cues elicited greater fMRI activity for valid than invalid cues in occipital areas (detailed statistics are presented in **Supplementary Table 3**). Critically, these effects were consistently observed across participants (**Figure 5**). Taken together, these results confirm both of our predictions.

**Figure 5.**
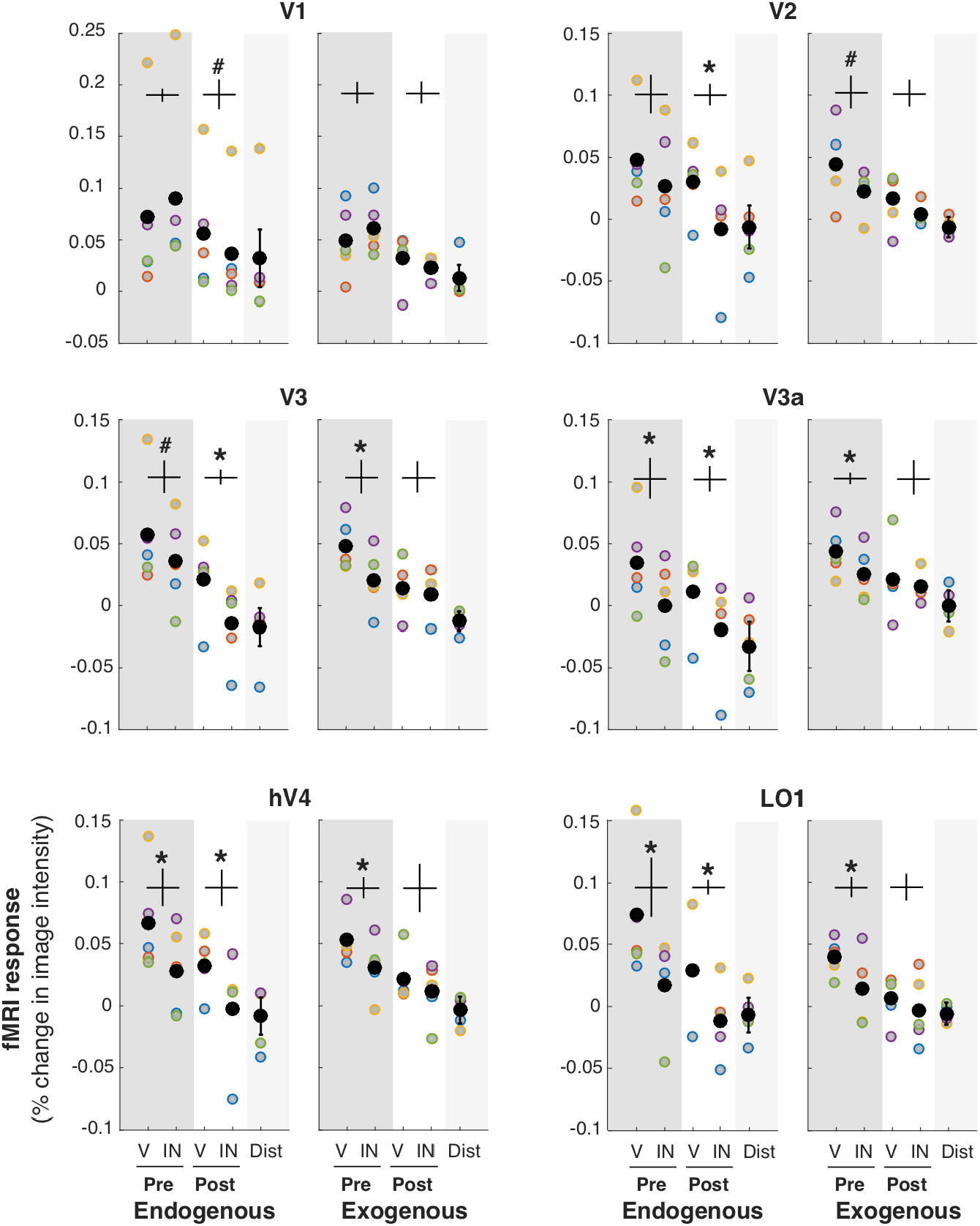
Specificity of each visual ROI for endogenous and exogenous attention. fMRI response amplitude was measured for each attentional condition. V: valid cue condition (target location matches the location indicated by pre-cue/post-cue). IN: invalid cue condition (target location at the opposite location relative to the pre-cue/post-cue). Pre: pre-cue presented before the grating stimuli. Post: post-cue presented after gratings. All four conditions in the contralateral ROI. Distractor (Dist): pre-cue and post-cue valid and invalid conditions averaged in the ipsilateral ROI. Each colored point corresponds to the data of one participant. The black points represent the average across all 5 participants. *, Statistically significant difference between valid and invalid, separately for pre and post-cueing (p < 0.05). #, trend (p < 0.1). Detailed statistics are presented in **Supplementary Table 1**. Error bars (vertical lines) are plotted on the mdifference between valid and invalid and represent ± 1 SEM across participant.

## DISCUSSION

This is the first study to compare pre and post-orienting, and reorienting, of endogenous and exogenous attention in visual cortex, while concurrently assessing visual performance using well-established psychophysical protocols to manipulate attention. The fact that the same participants performed the same orientation discrimination task with the same stimuli and task difficulty under different attentional manipulations enabled us to isolate the fMRI activity induced by each type of attention during orienting and reorienting.

Previous studies comparing endogenous and exogenous attention conditions state various attentional effects in early visual areas (see **Table 1**). Some report differential effects between these attention conditions in the right LO and MT ^52^, others in the cuneus ^51–53^, the occipital gyrus (SOG, MOG, LOG; ^52,53,64^), the fusiform area ^52,64,65^, and the TOS, LG and SC ^64^. This lack of clear picture regarding the differential impact of endogenous and exogenous spatial attention is likely due to the methodological differences and limitations we discussed in the Introduction (see also **Table 1**). In any case, it is often assumed that the effects of endogenous and exogenous attention are the same in visual occipital areas ^33,38,39,50,51^.

Other regions in striate and extra-striate visual areas could be implicated in both endogenous and exogenous attention. For instance, area V6 has been reported to be involved in the shift of covert attention, in both monkeys ^101,102^ and humans ^103^. Because this area has a large representation of the periphery ^104^, we would have needed a wider field of view than the one we had to reliably define it. Similarly, higher-level visual areas are also involved in shifts of covert spatial attention, such as IPS 0 to IPS 5 (e.g. ^31,39^). However, given the goal of our study, and thus our slice prescription, we could not identify all IPS regions.

To further our knowledge of the neural correlates of attention, we investigated both attentional orienting (valid cueing) and reorienting (invalid cueing), critical in an ever-changing environment ^42,80,81,94^. Furthermore, given ubiquitous performance tradeoffs between attended (benefits) and unattended (costs) locations (e.g. ^3,28,88–93^), we assessed activity at both attended (contralateral ROI) and unattended (ipsilateral ROI) locations. Finally, we investigated how attentional effects varied as a function of pre- and post-cueing, thus contrasting the neural correlates of perceptual and post-perceptual processing of information.

The behavioral effects obtained in the scanner are consistent with psychophysical studies. The enhanced performance brought about by the valid, but uninformative peripheral precue is consistent with an automatic, bottom-up involuntary capture of exogenous attention (e.g. ^3,4,7,8,10,13,15,28,47,48,54,77,88,89,92,105,106^). The enhanced performance brought about by the valid, informative central precue is consistent with a top-down, voluntary deployment of endogenous attention (e.g. ^3–8,11–13,26,28,47,48,54,87,92,107,108^).

In the endogenous attention condition, there was an increase in attentional modulation of stimulus-evoked activity along the hierarchy of visual areas. There is no consensus regarding the visual hierarchy beyond area V3 (e.g. ^109–113^), especially regarding V3A. However, most authors agree that hV4 precedes LO1 ^109–112^. In any case, our data are consistent with either a strict hierarchy or with V3A and hV4 being at the same level. Such a pattern is consistent with previous studies suggesting that endogenous attention is a top-down modulation from frontal and parietal areas feeding back to visual cortex, with diminishing effects in earlier visual areas ^31,39,82,83^. Inconsistent with previous studies (e.g. ^26,28,32,114,115^), there was no evidence for attentional modulation in V1. It might be that attentional modulation of V1 activity is more variable than other visual cortical areas, making it harder to detect (see also ^31,77^). Methodological differences between this and previous studies may have contributed to weakening the effect of attention in V1. The accrual time in the current endogenous condition was relatively short (1300 ms in the valid condition and 500 ms in the invalid condition) compared to previous studies investigating endogenous, voluntary attention, in which the cue and/or stimuli were presented for a long duration to maximize BOLD measurements (e.g. ^26,28,32,114,115^). We chose the minimum SOA (317 ms) at which performance benefits plateau to have as comparable conditions as possible to the accrual time with the SOA for exogenous attention (117 ms). This short accrual time may have limited the effects of attentional feedback to V1.

The fact that we do not see attentional modulation in V1 with our protocol may indicate that V1 is not always necessary for attention to exert its behavioral effects. However, this null result should not be over-interpreted so as to accept the null hypothesis. For example, the inconsistent findings in the EEG literature regarding whether spatial attention can modulate V1 activity during the initial wave of visual processing depends on several factors (e.g., location, presence of distractors, attentional load ^116^). Similarly, some fMRI papers have shown significant modulation of spatial attention in V1 (e.g. ^26,28,32,114,115^). Furthermore, a recent study has shown that TMS on the occipital pole (V1/V2) after attentional deployment eliminated the typical benefits and costs brought about by exogenous attention at the respective attended and unattended locations ^89^.

In the exogenous attention condition, in contrast to the endogenous attention, the attentional modulation was approximately constant across the visual hierarchy. Some previous studies have reported a similar effect ^117,118^, others a decrease ^119^, and yet others an increase ^77,120^ across the visual areas. This difference might be explained by different task parameters. For example, in the Liu et al. (2005) study ^77^, participants knew which of the two stimuli was the target they had to discriminate while the stimuli were presented, as one stimulus was vertical and the other was tilted to the left or the right. In the present study, both stimuli were independently tilted and participants had to wait for the response cue to appear to know which one was the target and which one was the distractor.

Unlike in the endogenous pre-cueing condition in which the attention effect increased along the processing stream, for the endogenous post-cueing effect there was no evidence that the attentional modulation varied across these visual areas. The constant effect in the post-cue condition could be due to the contribution of two counteracting factors: (1) the fMRI response evoked by the stimulus in early visual areas may decrease along the visual hierarchy ^121^; (2) the top-down modulations from frontal and parietal areas feedback to visual cortex with diminishing effects in earlier visual areas ^26,31,39,82,83,122^.

In the exogenous condition, there was no significant post-cueing effect on early visual areas. This result is consistent with that of Liu et al. (2005) ^77^, who while evaluating exogenous attention effects on occipital cortex included a post-cue condition to rule out sensory contamination of the cue (i.e. sensory response evoked by the cue itself) contributing to the enhanced BOLD activity found in their pre-cue condition. In addition to ruling out a possible sensory contamination, the present results show that, in contrast to endogenous attention, exogenous attention does not aid in the selective readout of information. Although there was no reliable modulation of exogenous post-cue on early and intermediate visual areas, in this study, it had an effect on behavior. Such effect is sometimes ^71,72^ but not always ^73–76^ observed. It is thus possible that our imaging protocol was not sensitive enough or that this behavioral effect is supported by higher order regions.

The ROI-based analysis that we followed here enabled us to compare contralateral and ipsilateral modulation of BOLD activity, thus providing additional information regarding the differences in processing dynamics of both types of attention. We observed a larger difference between contralateral and ipsilateral areas for the valid than the invalid cueing condition. This effect could be due to the fact that for the former, participants were attending to the same location throughout the trial, whereas for the latter, when the response cue did not match the precue, participants had to switch their spatial attention to the opposite stimulus location, thus activity at that new location would be accumulated for less time. For instance, for endogenous attention, for the valid condition participants could have been processing the target for almost 500 ms before the response cue appeared. When the response cue matched the pre-cue, participants continued processing and reading out the signal from that location for up to 800 ms (they were not allowed to give an answer before the end of the response cue period). But when the response cue did not match the pre-cue, then participants had to switch after 500 ms to the other location (ipsilateral) thus accumulating less activity. Similarly, the accumulation time for the invalid cue condition in exogenous attention was only about 300 ms. This accrual time explanation could also account for the larger difference between contralateral and ipsilateral for pre-cues than post-cues, i.e. there is a 300 ms accumulation when the exogenous pre-cue is invalid, while only 100 ms when the post-cue is invalid. Likewise, the larger modulatory effect for endogenous than exogenous attention is consistent with the difference in accrual time.

In a recent study ^42^ we demonstrated that sub-regions of the Temporo-Parietal Junction (TPJ) that respond specifically to visual stimuli are more active when attention needs to be spatially reoriented (invalid cueing) than when attention remains at the cued location (valid cueing), and that partially overlapping specific visual sub-regions mediate such reorienting of endogenous or exogenous attention. Both the present and the TPJ studies provide a comprehensive investigation of endogenous and exogenous attention, and pave the way for rigorous psychophysics informed, neuroimaging studies of covert, spatial attention. Here, we concentrated the analysis on visual cortical areas in the occipital lobe. The present findings further our knowledge of the neurophysiological bases of covert attention and have implications for models of visual attention, which should consider not only the similarities, but also the differences in the orienting and reorienting of endogenous and exogenous attention in occipital areas reported here.

## CONCLUSION

The present results show some similarities and reveal important differences in the specific neural correlates of endogenous and exogenous attention on early vision: An increasing modulation of fMRI activity for pre-cueing endogenous attention, but constant modulation for exogenous attention, along the hierarchy of visual occipital areas, as well as a reliable and constant modulation of fMRI activity for post-cueing endogenous attention in occipital areas but not for exogenous attention. These results suggest that endogenous attention facilitates both the encoding and the readout of visual information whereas exogenous attention only facilitates the encoding of information.

## MATERIALS and METHODS

The behavioral methods employed in this study and the behavioral results are the same as those we reported in a recent study, in which we compared activity in TPJ during orienting and reorienting of endogenous and exogenous attention ^42^. To maximize the effects of these two types of attention, i.e. the benefits at the attended location and concurrent costs at the unattended location, we used optimal spatial and temporal parameters (reviews by ^1,2^). To enable direct comparison between endogenous and exogenous attention, the same participants performed the same orientation discrimination task under both types of attention. The fMRI methods employed in this study are the same as those used in that study ^42^, but here, instead of analyzing TPJ activity, we analyzed activity in occipital areas.

### Participants

Five participants (two male and three female, 24-30 years-old) participated in the study. They all had normal or corrected-to-normal vision. The University Committee on Activities Involving Human Subjects at New York University approved the experimental protocol (IRB # 10-7094), and participants provided written informed consent. All methods were performed in accordance with US regulations and the Declaration of Helsinki. Our study used single-participant ROI-based analysis, and thus had a small sample size. The same sample size that has been used in many fMRI studies in our labs (e.g. ^26,28,42,77,123–127^), as well as in other labs (e.g. ^128–132^). Each participant performed nine scanning sessions: one session to obtain a set of three high-resolution anatomical volumes, two sessions for retinotopic mapping, three sessions for the exogenous attention condition and three sessions for the endogenous attention condition (with the order counterbalanced among participants). Participants performed several practice sessions outside the scanner prior to the first scanning session of each attention condition.

### Stimuli

Stimuli were generated on a Macintosh computer using the MGL toolbox ^133^ in MATLAB (MathWorks). Stimuli were presented on a flat-panel display (NEC, LC-XG250 MultiSync LCD 2110; refresh rate: 60 Hz; resolution: 1024 × 768 pixels) positioned at the rear of the scanner bore and housed in a Faraday box with an electrically conductive glass front. The display, calibrated and gamma corrected using a linearized lookup table, was at a viewing distance of 172 cm from the participant, and visible through an angled mirror attached to the head coil. A central, white fixation cross (0.3°) was presented throughout the experiment. The two stimuli were two 4-cpd gratings windowed by raised cosines (3° of diameter; 7% contrast), one in each bottom quadrant (5° horizontal eccentricity; –2.65° altitude; 5.66° of eccentricity from the central fixation to the stimulus center). Both endogenous cues and exogenous cues were white rectangles (0.7°). The endogenous cues appeared adjacent to the fixation cross indicating one of the two lower quadrants (0.35° horizontal eccentricity from the edge of the fixation cross, and 0.35° altitude). The exogenous cues appeared adjacent to an upcoming grating stimulus, vertically aligned with the stimulus and above the horizontal meridian (1° away from the edge of the grating stimulus; and the edge of the cue 4.44° horizontal eccentricity from the edge of the fixation cross).

### Behavioral procedure

An exogenous attention condition trial lasted 1700 ms, whereas an endogenous attention condition trial lasted 1900 ms, the only difference being the stimulus-onset asynchronies (SOA) between the cue and the display; the timing of all the visual stimuli was the same in both attention conditions (**Figure 1**; the display is not at scale for illustration purposes). In the precue condition (40% of the trials), a cue preceded the two gratings. In 40% the post-cue condition, the cue followed the presentation of the gratings. In ‘cue-only’ trials (10% of the trials), the gratings were not presented. In ‘blank’ trials (10% of the trials), neither a cue nor the gratings were presented. These trials were then included in the GLM analysis to model the contribution of the visual signal produced by the cue (see MRI procedure). For both pre-cue and post-cue trials, participants were asked to press one of two keys to report the orientation of a target grating, i.e., clockwise or counter-clockwise compared to vertical. Participants pressed a third key in the case of cue-only and blank trials.

In both exogenous and endogenous condition, cues were presented for 67 ms, indicating either the bottom left or right quadrant of the screen. The inter-stimulus interval (ISI) between the cue and the grating stimuli was 50 ms for exogenous and 250 ms for endogenous conditions, resulting in SOA of 117 ms and 317 ms. We used the same timings for pre- and post-cue conditions (e.g. ^10,26,66,77^). These delays are optimal to manipulate exogenous and endogenous attention, while keeping the trial duration as similar as possible, and have been shown to maximize the behavioral consequences of each attention condition ^47,134–136^.

The behavioral effects of endogenous attention are sustained (e.g. ^107^) and thus, as shown in ERP studies (e.g. ^137^), are still present in later brain activity. Additionally, during 300 ms following cue onset, the brain responses elicited by exogenous and endogenous cues differ (for review ^1^). The two grating stimuli were then displayed for 50 ms. For the post-cue trials we kept the timings of cue and stimuli constant but inverted the order of their presentation (e.g.^10,26,66,77^). A response cue, presented for 800 ms at the end of the trial after both the cue and the stimuli had disappeared, indicated which one of the two gratings was the target (50% of the trials on the right and the remaining 50% and on the left). The maximum delay between the offset of the grating stimuli and the onset of the response cue was shorter (~400 ms max in the endogenous condition) than typically associated with a demand for working memory (>600 ms; ^138^). Immediately following each trial, a change of color of the fixation cross provided visual feedback to the participants, i.e. green for correct or red for incorrect responses. The fixation cross did not change color if participants had missed the response window, i.e. if they had not pressed any key after 530 ms.

In the exogenous attention condition, a peripheral cue was presented, which was not informative regarding the target location or orientation. When the cue location matched the target location, it was considered a valid trial (50% of the trials), otherwise it was considered an invalid trial (the remaining 50% of the trials). In the endogenous attention condition, a central cue pointed to either the left or right quadrant. The cue was informative of the target location but not its orientation (75% valid trials and 25% invalid trials). Participants were informed of this validity. It is important to note that: (1) cue validity does not affect cueing effectiveness for exogenous attention, although it does so for endogenous attention (e.g. ^3,139,140^); (2) the short timing and non-informative cue in the exogenous condition ensured that no voluntary component could be involved in the exogenous attentional effect.

Endogenous and exogenous attention conditions were performed in separate sessions to ensure optimal manipulation of each attention system. Participants performed two practice sessions outside the scanner before the first session of each attentional scanning condition. To equate task difficulty for both attention conditions, using a staircase procedure, the tilt of the target grating was adjusted for each participant to achieve ~80% correct performance in the valid trials in each attention condition. In each of the six experimental scanning sessions (three sessions of exogenous attention and three of endogenous), participants performed 14 runs of 40 trials each, as well as a run of stimulus localizer (see MRI procedure). The tilt was then adjusted between runs to maintain overall performance at ~80% correct.

Eye position was monitored during all scanning sessions using an infrared video camera system (Eyelink 1K, SR Research, Ottawa, Ontario, http://www.sr-research.com/EL_1000.html). Trials in which the participants blinked or broke fixation (1.5° radius from central fixation) at any point from fixation onset to response cue offset were identified and regressed separately in the MRI analysis, and removed from the behavioral analysis (13% ± 4% of the trials on average across all participants).

### MRI Procedure

#### Scanning

Imaging was conducted on a 3T Siemens Allegra head-only scanner (Erlangen, Germany), using a Siemens NM-011 head coil (to transmit and receive) to acquire anatomical images, a receive-only 8-channel surface coil array (Nova Medical, Wilmington, MA) to acquire functional images. To minimize participants’ head movements, padding was used.

For each participant, three high-resolution anatomic images were acquired in one scanning session, using a T1-weighted magnetization-prepared rapid gradient echo (MPRAGE) sequence (FOV = 256 × 256 mm; 176 sagittal slices; 1 × 1 × 1 mm voxels), and were coregistered and averaged. Using FreeSurfer (public domain software; http://surfer.nmr.mgh.harvard.edu), the gray matter was segmented from these averaged anatomical volumes. All subsequent analyses were constrained only to voxels that intersected gray matter.

T2*-weighted echo-planar imaging sequence (i.e. functional images; TR = 1750 ms; TE = 30 ms; flip angle = 90°) measured blood oxygen level-dependent (BOLD) changes in image intensity ^141^. In each volume, 28 slices covered the occipital and posterior parietal lobes and were oriented 45° to the calcarine sulcus (FOV = 192 × 192 mm; resolution = 2 × 2 × 2.5 mm; no gap). To align functional images from different sessions to the same high-resolution anatomical volume for each participant, we acquired an additional T1-weighted anatomical images in the same slices as the functional images (spin echo; TR = 600 ms; TE = 9.1 ms; flip angle = 90°; resolution = 1.5 × 1.5 × 3 mm) during each scanning session, and used them in a robust image registration algorithm.

#### MRI data pre-processing

Imaging data were analyzed in MATLAB, using mrTools ^142^ and custom software. To allow longitudinal magnetization to reach steady state, the first eight volumes of each run were discarded. Spatial distortion was corrected using the B0 static magnetic field measurements performed in each session. The functional data were then motion corrected, the linear trend was removed, and a temporal high-pass filter was applied (cutoff: 0.01 Hz) to remove slow drifts and low-frequency noise in the fMRI signal.

#### Retinotopic mapping

We followed well-established conventional traveling-wave, phase-encoded methods. Using clockwise and counter-clockwise rotating checkerboard wedges, we measured phase maps of polar angle. Using contracting and expending checkerboard rings, we measured eccentricity maps ^95–98^. **Figure 2**(left panel) shows the visual areas of interest that were drawn by hand on flattened surface of the brain, following published conventions ^97,98,124,143^.

#### Stimulus localizer

In each scanning session of the main experiment, participants completed one stimulus localizer run (6 runs overall, 4 min each). A run consisted of 16 cycles (17.5 s) of a block alternation protocol between stimulus on (8.75 s) and stimulus off (8.75 s). Participants only had to fixate the central cross throughout each run. The stimuli were at the same location, and of the same size and spatial frequency as those in the main experiment, except at full contrast and their phase and orientation changed randomly every 200 ms to avoid adaptation. To define the cortical representation of the gratings, we then averaged the data across the 6 runs and followed the same methods as for the retinotopic mapping. Voxels that responded positively during the blocks when the grating stimuli were presented were used to restrict each retinotopic ROI. The fMRI time series from each voxel were fit to a sinusoid. To be conservative, only voxels whose best-fit sinusoid had a phase value between 0 and pi, and a coherence between the best-fit sinusoid and the time series greater than 0.2 were included in the ROI (**Figure 2**, right panel). Analysis performed without restricting the ROI to this coherence level yielded similar results.

#### Event-related analysis

fMRI time series were averaged across voxels in each ROI (separately for each hemisphere) and then concatenated across runs. The data were denoised using GLMDenoise ^144^, and fMRI response amplitudes were computed using linear regression, with twelve regressors: 8 combinations of right and left valid and invalid pre- and post-cue, right and left cue-only, blank (no cue nor stimulus) and eye-movements (blink or broken fixation). For each ROI in each hemisphere, the resulting fMRI response amplitudes (for correct trials only) were then averaged across participants.

## Acknowledgments

This work was supported by: NIH RO1-EY019693 to MC and DJH; NIH RO1-EY027401 to MC; the FYSSEN Foundation and the Philippe Foundation to LD; and the Center for Brain Imaging of New York University. We want to thank members of the Carrasco lab, especially Rachel Denison and Michael Jigo, as well as Stephanie Badde and Jonathan Winawer for constructive comments on the manuscript.

## Competing interests

The authors declare no competing interests.

## Author contributions

### Laura Dugué

Conceptualization, Data curation, Formal analysis, Funding acquisition, Investigation, Visualization, Methodology, Writing - original draft, Writing - review and editing.

### Elisha Merriam

Data curation, Formal analysis, Investigation, Visualization, Methodology, Writing - review and editing.

### David Heeger

Resources, Supervision, Funding acquisition, Visualization, Methodology, Writing - review and editing.

### Marisa Carrasco

Conceptualization, Resources, Supervision, Funding acquisition, Visualization, Methodology, Writing - original draft, Project administration, Writing - review and editing.

## Data availability

Data relative to the main results of this study (i.e. Figures 3, 4 and 5) are made available via OSF at [link provided upon acceptance of the manuscript].

## Notes

### Competing Interest Statement

The authors have declared no competing interest.

