## Supplementary information for "Differential impact of endogenous and exogenous attention on activity in human visual cortex"

–

**Running title:** endogenous and exogenous attention

**Corresponding author:**
Laura Dugué
Current address: 45 rue des Saints-Pères 75006 Paris, FRANCE


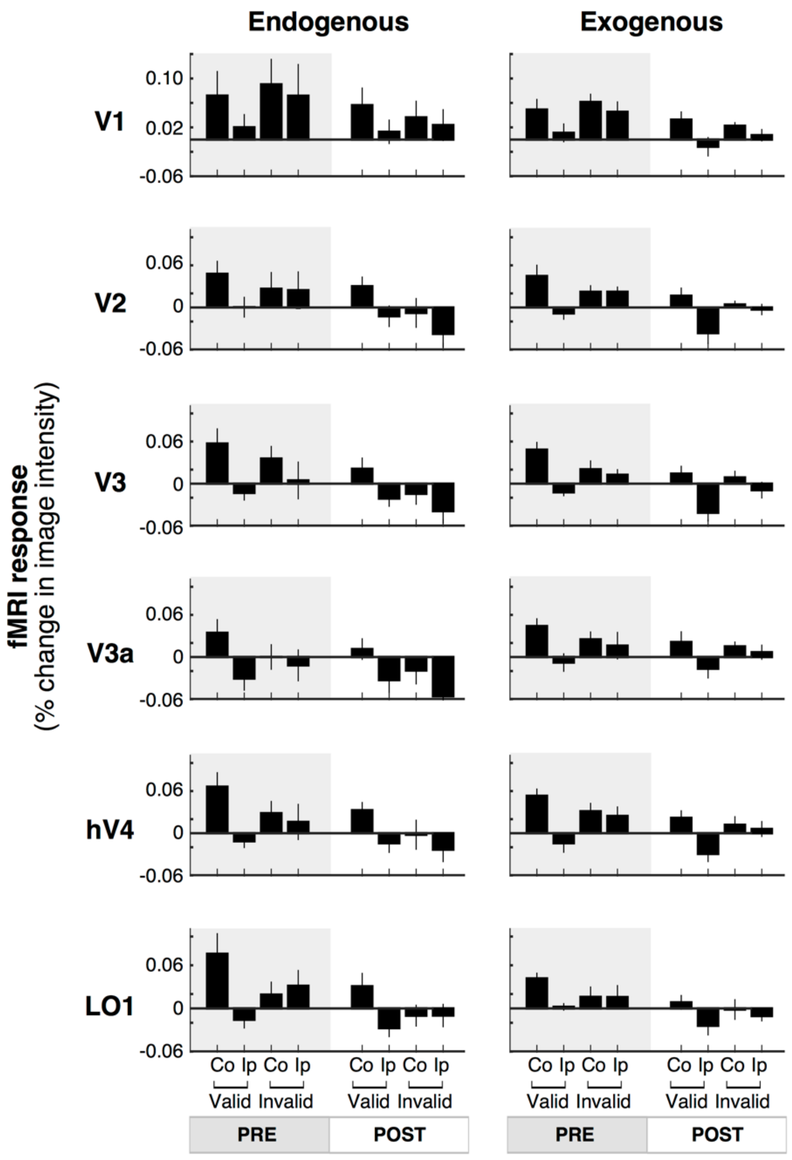


***Supplementary Figure 1.*** *Endogenous and exogenous attention effects in each contralateral and ipsilateral visual ROI. fMRI response amplitude was measured for each attentional condition, separately in contralateral (Co) and ipsilateral (Ip) ROIs relative to the cued location. V: valid cue condition (target location matches the location indicated by pre-cue/post-cue). IN: invalid cue condition (target location at the opposite location relative to the pre-cue/post-cue). Pre: pre-cue presented before the grating stimuli. Post: post-cue presented after grating stimuli. For both attention types, contralateral activity was higher than ipsilateral activity and this difference was more pronounced for valid than invalid trials. Error bars represent ± 1 SEM.*

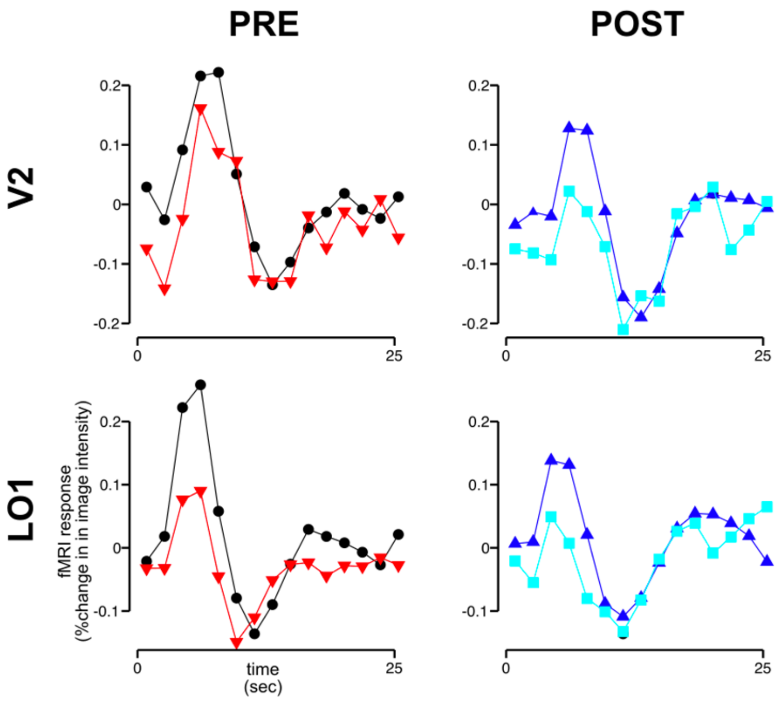


***Supplementary Figure 2. fMRI time courses for endogenous attention for a representative participant, two specific ROIs.*** *Black, valid pre-cue condition. Red, invalid pre-cue. Blue, valid post-cue. Cyan, invalid post-cue.*

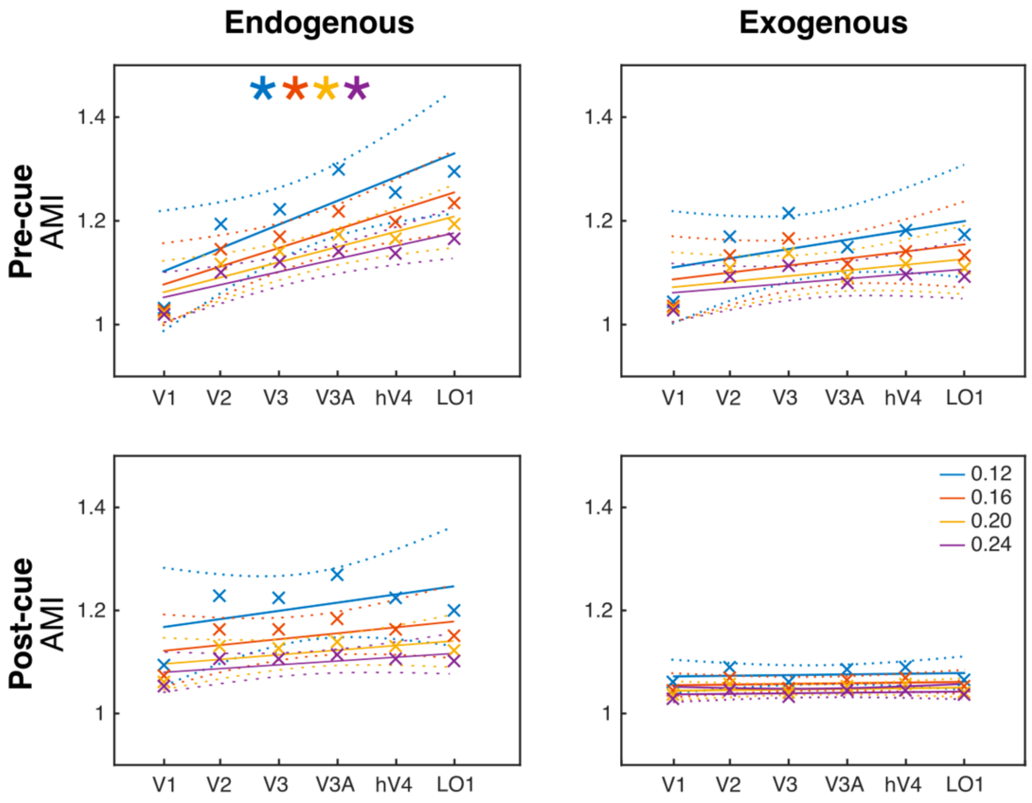


***Supplementary Figure 3. Regression analyses relative to Figure 3 performed for 4 different constant values (colors).*** AMI, *Attentional Modulation Index separately for pre- and post-cue conditions for each ROI (see main manuscript). *, Statistically significant regression analysis ($p < 0.05$).*

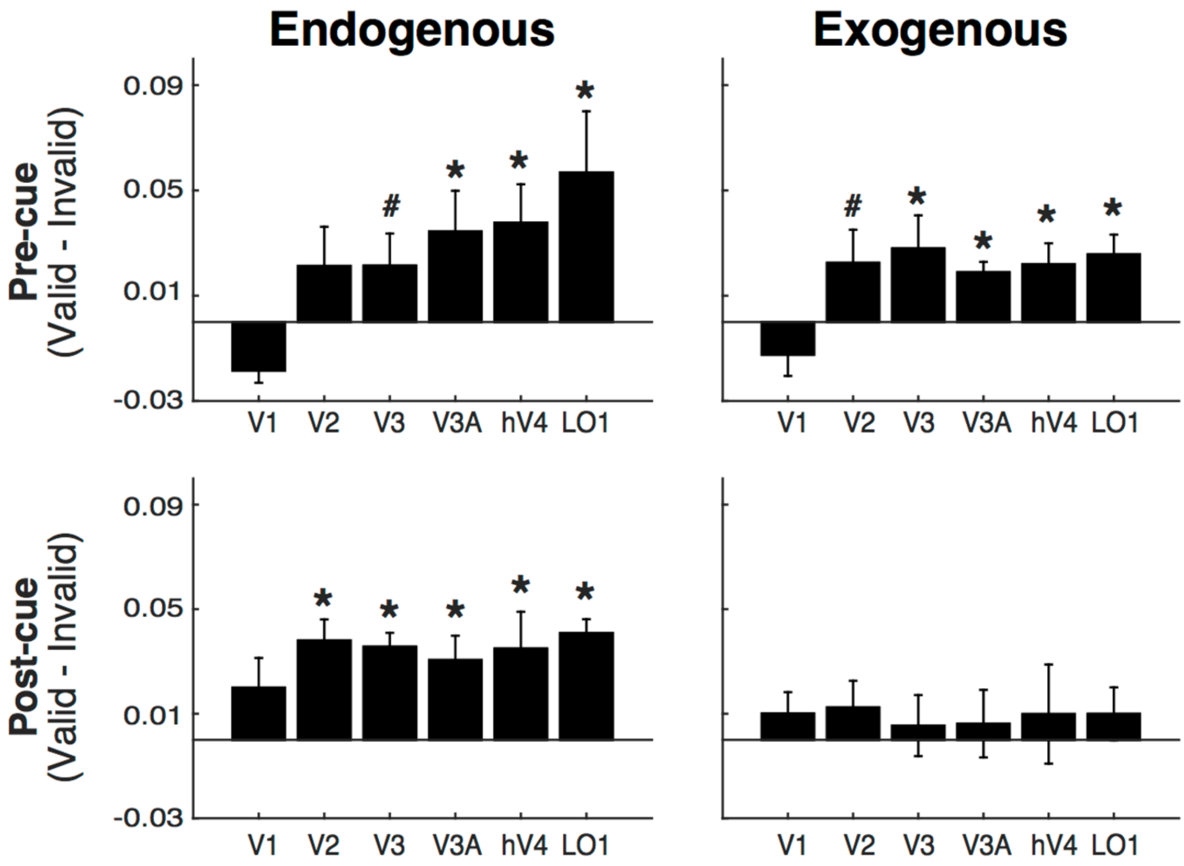


***Supplementary Figure 4.*** **Single ROI responses (percent signal change) for pre *and post-cueing and for both endogenous and exogenous attention*** *(relative to Figure 4). The difference between valid and invalid conditions is plotted separately for pre- and post-cue conditions for each ROI. *, Statistically significant difference between valid and invalid ($p < 0.05$). Error bars on plots are ± 1 SEM.*

| | PRE | | | | POST | | | |
|---|---|---|---|---|---|---|---|---|
| **Constant value** | **0.12** | **0.16** | **0.20** | **0.24** | **0.12** | **0.16** | **0.20** | **0.24** |
| Intercept | t(56)=8.6, **p<0.001**, e=1.0±0.1 | t(56)=11.0, **p<0.001**, e=1.0±0.09 | t(56)=13.2, **p<0.001**, e=1.0±0.08 | t(56)=15.4, **p<0.001**, e=1.0±0.07 | t(56)=13.5, **p<0.001**, e=1.2±0.09 | t(56)=18.7, **p<0.001**, e=1.2±0.06 | t(56)=23.3, **p<0.001**, e=1.1±0.05 | t(56)=27.8, **p<0.001**, e=1.1±0.04 |
| ROI | t(56)=3.2, **p=0.002**, e=0.07±0.02 | t(56)=3.4, **p=0.001**, e=0.06±0.02 | t(56)=3.5, **p=0.001**, e=0.05±0.01 | t(56)=3.6, **p=0.001**, e=0.04±0.01 | t(56)=1.4, p=0.16, e=0.03±0.02 | t(56)=1.5, p=0.13, e=0.02±0.02 | t(56)=1.6, p=0.13, e=0.02±0.01 | t(56)=1.6, p=0.12, e=0.02±0.01 |
| Endo/Exo | t(56)=0.6, p=0.57, e=0.04±0.06 | t(56)=0.7, p=0.49, e=0.03±0.05 | t(56)=0.7, p=0.46, e=0.03±0.04 | t(56)=0.8, p=0.44, e=0.02±0.03 | t(56)=-1.2, p=0.23, e=-0.07±0.06 | t(56)=-1.1, p=0.27, e=-0.05±0.04 | t(56)=-1.1, p=0.29, e=-0.04±0.03 | t(56)=-1.0, p=0.31, e=-0.03±0.03 |
| ROI x Endo/Exo | t(56)=-2.0, **p=0.046**, e=-0.03±0.01 | t(56)=-2.2, **p=0.032**, e=-0.02±0.01 | t(56)=-2.3, **p=0.025**, e=-0.02±0.01 | t(56)=-2.4, **p=0.021**, e=-0.02±0.01 | t(56)=-1.2, p=0.25, e=-0.02±0.02 | t(56)=-1.3, p=0.21, e=-0.01±0.01 | t(56)=-1.3, p=0.2, e=-0.01±0.01 | t(56)=-1.3, p=0.19, e=-0.01±0.01 |

***Supplementary Table 1. Statistics from LME performed for 4 different constant values.*** *A fixed constant was added to all condition values of each participant before computing the AMI so that all resulting values were positive. To ensure the results of the LME (see main manuscript) were not dependent on the value of this constant, the LMEs were performed on four different constant values. The results remain the same. In the manuscript we plot the data (**Figure 3**) and report the statistics for constant value = 0.2.*

| | Endogenous | | | | Exogenous | | | |
|---|---|---|---|---|---|---|---|---|
| **Constant value** | **0.12** | **0.16** | **0.20** | **0.24** | **0.12** | **0.16** | **0.20** | **0.24** |
| **Pre** | F(4)=10.9, **p=0.03**, $R^2$=0.73 | F(4)=14.1, **p=0.02**, $R^2$=0.78 | F(4)=16.5, **p=0.02**, $R^2$=0.81 | F(4)=18.4, **p=0.01**, $R^2$=0.82 | F(4)=1.9, p=0.24, $R^2$=0.32 | F(4)=1.9, p=0.25, $R^2$=0.32 | F(4)=1.8, p=0.25, $R^2$=0.31 | F(4)=1.8, p=0.25, $R^2$=0.31 |
| **Post** | F(4)=1.3, p=0.31, $R^2$=0.25 | F(4)=1.8, p=0.25, $R^2$=0.31 | F(4)=2.2, p=0.22, $R^2$=0.35 | F(4)=2.4, p=0.20, $R^2$=0.38 | F(4)=0.1, p=0.75, $R^2$=0.03 | F(4)=0.2, p=0.67, $R^2$=0.05 | F(4)=0.3, p=0.61, $R^2$=0.07 | F(4)=0.4, p=0.57, $R^2$=0.09 |

***Supplementary Table 2. Statistics from linear regressions performed for 4 different constant values.*** *The results remain the same. In the manuscript we plot the data (**Figure 3**) and report the statistics for constant value = 0.2.*

| | Endogenous | | | | | | Exogenous | | | | | |
|---|---|---|---|---|---|---|---|---|---|---|---|---|
| | V1 | V2 | V3 | V3A | hV4 | LO1 | V1 | V2 | V3 | V3A | hV4 | LO1 |
| **Pre** | t(4)=1.3, p=0.13, CI=0.99 | t(4)=3.0, **p=0.02**, CI=1.03 | t(4)=4.2, **p=0.007**, CI=1.07 | t(4)=3.4, **p=0.014**, CI=1.07 | t(4)=4.1, **p=0.008**, CI=1.08 | t(4)=4.5, **p=0.005**, CI=1.1 | t(4)=1.2, p=0.157, CI=0.97 | t(4)=2.6, **p=0.03**, CI=1.02 | t(4)=3.7, **p=0.01**, CI=1.06 | t(4)=8.8, **p=0.0005**, CI=1.07 | t(4)=5.2, **p=0.003**, CI=1.07 | t(4)=9.0, **p=0.0004**, CI=1.08 |
| **Post** | t(4)=2.2, **p=0.047**, CI=1.00 | t(4)=4.3, **p=0.007**, CI=1.07 | t(4)=11.5, **p=0.0002**, CI=1.10 | t(4)=3.4, **p=0.013**, CI=1.05 | t(4)=3.7, **p=0.011**, CI=1.05 | t(4)=8.9, **p=0.0004**, CI=1.09 | t(4)=1.5, p=0.107, CI=0.98 | t(4)=1.8, p=0.074, CI=0.99 | t(4)=1.6, p=0.09, CI=0.98 | t(4)=1.0, p=0.198, CI=0.95 | t(4)=1.4, p=0.122, CI=0.97 | t(4)=1.8, p=0.077, CI=0.99 |

***Supplementary Table 3. Statistics from Figure 4 and 5.*** *One-tailed t-tests are computed on the AMI separately for pre and post-cue conditions for each ROI against a ratio of 1 (no effect). CI, Confidence Interval.*
